# A comprehensive catalog of 3D genome organization in diverse human genomes facilitates understanding of the impact of structural variation on chromatin structure

**DOI:** 10.1101/2023.05.15.540856

**Authors:** Chong Li, Marc Jan Bonder, Sabriya Syed, Human Genome Structural Variation Consortium (HGSVC), HGSVC Functional Analysis Working Group, Michael C. Zody, Mark J.P. Chaisson, Michael E Talkowski, Tobias Marschall, Jan O Korbel, Evan E Eichler, Charles Lee, Xinghua Shi

**Affiliations:** Temple University, Department of Computer and Information Sciences, Philadelphia, PA, USA; German Cancer Research Center (DKFZ), Division of Computational Genomics and Systems Genetics, Heidelberg, Germany; The Jackson Laboratory for Genomics Medicine, Farmington, CT, USA; New York Genome Center, New York, NY, USA; Department of Quantitative and Computational Biology, University of Southern California, Los Angeles, CA, USA; Program in Medical and Population Genetics, Broad Institute of MIT and Harvard, Cambridge, MA, USA; Center for Genomic Medicine, Massachusetts General Hospital, Boston, MA, USA; Department of Neurology, Massachusetts General Hospital and Harvard Medical School, Boston, MA, USA; Stanley Center for Psychiatric Research, Broad Institute of MIT and Harvard, Cambridge, MA, USA; Institute for Medical Biometry and Bioinformatics, Medical Faculty, Heinrich Heine University, Düsseldorf, Germany; Center for Digital Medicine, Heinrich Heine University, Düsseldorf, Germany; European Molecular Biology Laboratory (EMBL), Genome Biology Unit, Heidelberg, Germany; University of Washington School of Medicine, Department of Genome Sciences, Seattle, WA, USA; Howard Hughes Medical Institute, University of Washington, Seattle, WA, USA

**Keywords:** Hi-C, 3D genome, topologically associating domains, chromatin loops, structural variation, gene regulation, human genome

## Abstract

The human genome is packaged into the three-dimensional (3D) nucleus and organized into functional units known as topologically associating domains (TADs) and chromatin loops. Recent studies show that the 3D genome can be modified by genome structural variants (SVs) through disrupting higher-order chromatin organizations such as TADs, which play an essential role in insulating genes from aberrant regulation by regulatory elements outside TADs. Here, we have developed an integrative Hi-C analysis pipeline to generate a comprehensive catalog of TADs, TAD boundaries, and loops in human genomes to fill the gap of limited resources. We identified 2,293 TADs and 6,810 sub-TADs missing in the previously released TADs of GM12878. We then quantified the impact of SVs overlapping with TAD boundaries and observed that two SVs could significantly alter chromatin architecture leading to abnormal expression and splicing of genes associated with human diseases.

## Introduction

To better understand the genotype-to-phenotype relationship in the human genome, functional annotation of genomes is critical. As an intermediate step, it is vital to determine how the spatial arrangement of DNA impacts genome functionality and gene regulation through the characterization of the three-dimensional (3D) structure of chromatin inside the nucleus^14^. Chromosome conformation capture sequencing (Hi-C) is a genome-wide sequencing technique that combines proximity-based DNA ligation with high-throughput sequencing to measure the geographical proximity of possibly any pair of genomic loci^1^. Techniques such as Hi-C have been widely used to characterize the 3D structure of the genome and uncover folding principles of chromatin, including topologically associating domains (TADs) and chromatin loops. TADs are stable genomic regions separated by insulating proteins, e.g., CCCTC-binding factor (CTCF), and provide an encapsulating domain for constraining the chromatin contacts between regulatory elements and genes^2, 28, 29, 79^. TAD boundaries, which separate adjacent TADs, have been found to be well conserved across cell types and more evolutionarily constrained than TADs themselves^40^. At kilobase to megabase scale, Hi-C data have been utilized to pinpoint specific interactions between distant chromatin regions, known as chromatin loops, which are detected as enriched regions compared to their surrounding areas^52^. Chromatin loops typically connect promoters and enhancers and correlate with the activation of genes. It is shown that chromatin loops are conserved across different cell types and species^5^, and the rewiring of such loops contributes to developmental disorders and tumorigenesis^13, 53^.

Recently, there has been a growing interest in uncovering the disruption of gene regulation resulting from pathogenic large genomic rearrangements or structural variants (SVs), including deletions, insertions, inversions, and duplications. These genomic rearrangements can disturb the normal 3D structure of the genome and lead to aberrant interactions between chromatin-regulatory elements^44^, with such alterations bringing about the abnormal expression of oncogenic and disease-causing genes^42^. Studies have shown that disruptions to TADs can change the long-range regulatory architecture, result in alterations of gene expression levels, and lead to diseases^32^. In particular, SVs can affect CTCF-associated border elements, alter gene expression and enhancer-promoter interactions, and eventually result in human diseases^32, 34^. Another study found that SVs can cause distinct TADs to fuse, and complex rearrangements significantly alter the chromatin folding maps in cancer genomes^44^. Additionally, by integrating the information of TADs, the gene expression data, and somatic copy-number alterations (SCNAs), it can help to identify the cancer-related gene overexpression resulting from *cis*-regulatory elements reorganization, such as enhancer hijacking^43^. A recent study identified SV-induced enhancer-hijacking and silencer-hijacking events and reported the role of repressive loops on their target genes in patients with acute myeloid leukemia^51^.

Hence, it is of great need to characterize the 3D genome organization of human genomes to facilitate the understanding of the 3D structure landscape of human genomes and the impact of genetic variants on chromatin structures^66^. In doing this, Hi-C has been demonstrated as a powerful tool to chart the landscape of chromatin structures, including TADs and chromatin loops at the genome level^2^. To date, the most comprehensive characterization of 3D genome organization from Hi-C sequencing comes from GM12878, with the densest Hi-C data so far^5^, and individuals from the 4D Nucleome (4DN) project^24^. Despite these efforts, the utility of Hi-C data is typically limited due to their relatively low resolution, insufficient read coverage, and the sparse contact matrices that result in the lack of detailed characterization of chromatin structures in human genomes. Given the challenges in producing high-coverage Hi-C data due to sequencing costs and library preparation, various computational methods have been proposed to augment or enhance the quality of Hi-C data. For example, HiCPlus^45^ applies a straightforward neural network optimized using mean squared error, and an improved tool named HiCNN^46^ was proposed to improve Hi-C data quality. The use of Generative Adversarial Networks (GAN) was then introduced in the hicGAN^47^, which produces high-resolution contact maps based on low-resolution versions of such contact maps from Hi-C data. Another GAN-based method, DeepHiC^48^, was proposed with a user-friendly web interface. In HiCSR^49^, a novel loss function is tailored to optimize an adversarial loss in a GAN to improve GAN-based Hi-C data augmentation. Another method, VEHiCLE^50^, was presented to enhance Hi-C contact maps utilizing variational autoencoder generative models incorporating multiple loss functions.

These observations and studies highlight the importance of providing a comprehensive map of 3D chromatin structure to represent the spatial genomic organization in diverse human genomes in healthy populations. This comprehensive map in humans will facilitate the investigation of genomic variation and the disrupted 3D genome structures in health and disease. Therefore, in this study, we collected Hi-C data on the 1000 Genomes individuals^6^ from various projects to construct a comprehensive resource of TADs and loops in lymphoblastoid cell lines (LCLs) in 44 different individuals from five super populations including Africa (AFR), East Asia (EAS), Europe (EUR), South Asia (SAS), and the Americas (AMR) (Table S12). This Hi-C data set includes Hi-C sequencing of 27 individuals from HGSVC2^4^, six individuals for three trios from HGSVC1^22^, ten individuals from 4DN project^22, 24^, and GM12878^5^. We derived the TAD map of these samples by integrating two TAD/TAD boundary-calling methods, *Arrowhead*^5^ and *Insulation Score*^8^ *(IS),* to take advantage of their respective strengths. *Arrowhead* is able to call TADs that exhibit TAD-associated biological features and can output hierarchical TADs^58^. Meanwhile, TADs detected by *IS* are demonstrated to be robust against low coverage levels and capable of detecting dynamics of TAD boundary strength under different conditions^59^. We also generated a comprehensive catalog of chromatin loops in these individuals by combining loops produced from two different loop callers (*HiCCUPs*^5^ and *cooltools*^12^).

To demonstrate the utility of this map of 3D chromatin structure, we utilized the up-to-date genetic variation, particularly SVs of all major types, on the 64 haplotypes from 32 diverse human genomes produced by the Human Genome Structural Variation Consortium (HGSVC)^4^. Equipped with the comprehensive catalog of TADs and TAD boundaries resulting from this study, we assessed the impact of SVs on chromatin organization in a variety of human genomes that come from five super populations in the 1000 Genomes samples^6^. First, we identified 11,653 SVs that intersect with TAD boundaries with the possibility of disrupting local chromatin structures. Among these, two SVs, including one deletion (chr8-644401-DEL-5014) and one insertion (chr6-130293639-INS-295), were revealed to significantly alter TAD boundaries and also gene expression and splicing by overlapping with eQTL and sQTL signals previously reported on these samples^4^. Except for TADs, we then conducted individual loop calling and generated a comprehensive catalog of loops on the same samples. Our analysis illustrates the utility of this up-to-date 3D genome structure in diverse humans as a resource to help elucidate the links between genetic variation and 3D genome structure, as well as their disruptive impact on gene regulation.

## Results

### 1. A comprehensive TAD catalog identified across diverse individuals in human genomes

We first sought to construct a 3D chromatin structure map of human genomes in LCLs by combining Hi-C data from 44 individuals and comparing the 3D chromatin structure features across these diverse individuals (Table 1). Hi-C data collected on these 44 individuals were pre-processed using the same pipeline to generate the final map, including characterization of TADs, TAD boundaries, and chromatin loops. By an integrative analysis of the Hi-C data from these 44 individuals, we identified 18,972 TADs in our Integrative Catalog using a customized pipeline that combines two TAD calling algorithms, namely *Arrowhead and Insulation Score (IS)*, for calling TADs and quantifying TAD boundaries (Table S2-S5). We compared the results between our Integrative Catalog and the previously published largest map of GM12878 human LCLs (Table S1, Figure S3, Figure S4)^5^. We found that our Integrative Catalog covers all of the TADs of GM12878 released by ENCODE^60^ (Fig. 1a). Specifically, our Integrative Catalog identified 2,293 novel TADs and 6,810 smaller sub-TADs^5, 27, 76^ within 3,605 meta-TADs of GM12878 (ENCODE), that were missing in the GM12878 (ENCODE) (Fig. 1b). Note that, for any of the data and results on GM12878 included in this study, we did not use the original results published from Rao et al.^5^, but used the results re-processed by ENCODE^60^ that mapped to hg38 as the reference genome (Methods). Explicitly, we considered two TADs shared when their regions are over 50% reciprocally overlapped. Thus, around 95.34% and 74.30% of the TADs released by ENCODE for GM12878 coincided with the TADs separately detected by *Arrowhead* and *IS* in our study, respectively. Approximately 66.43% of the TADs called by *Arrowhead* can also be detected by *IS* in our dataset (Figure S2). We observed that the relatively lower number of overlapped TADs between *Arrowhead* and *IS* is due to the fact that *Arrowhead* tends to call TADs in larger sizes than *IS*, which cannot be recognized when we calculate the reciprocal overlap once the size of the TAD called in *Arrowhead* is more than half of the potential overlapped TAD called by *IS* (Figure S1). For a more strict overlapping comparison, we found that over 97.5% of the TADs released by ENCODE for GM12878 are over 50% reciprocally overlapped with our Integrative Catalog call set (Figure S1). Since there were no released TAD boundary locations for GM12878, we only focused on comparing the sizes and numbers of the TADs in this study (Table S1).

**Fig. 1.**
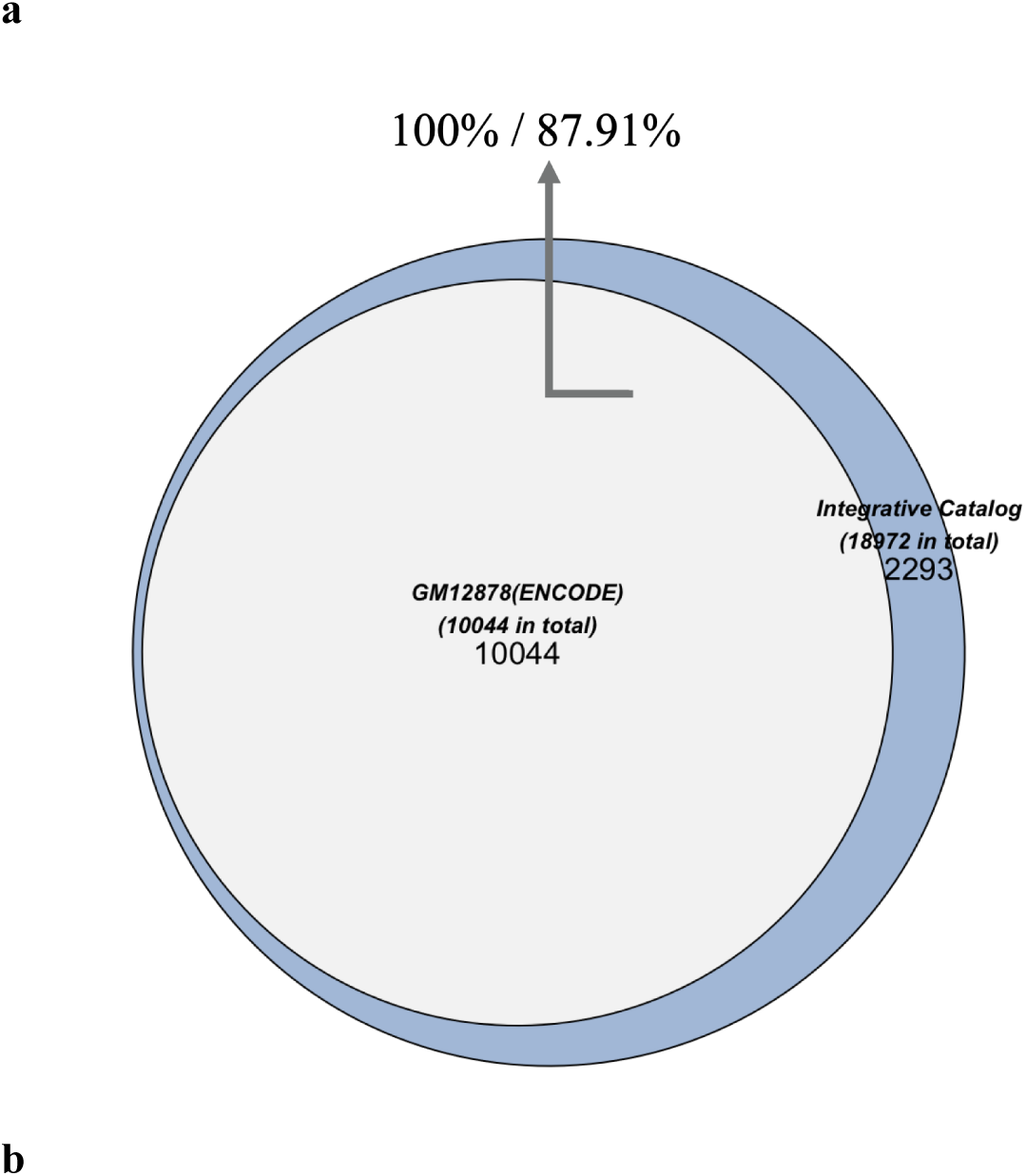
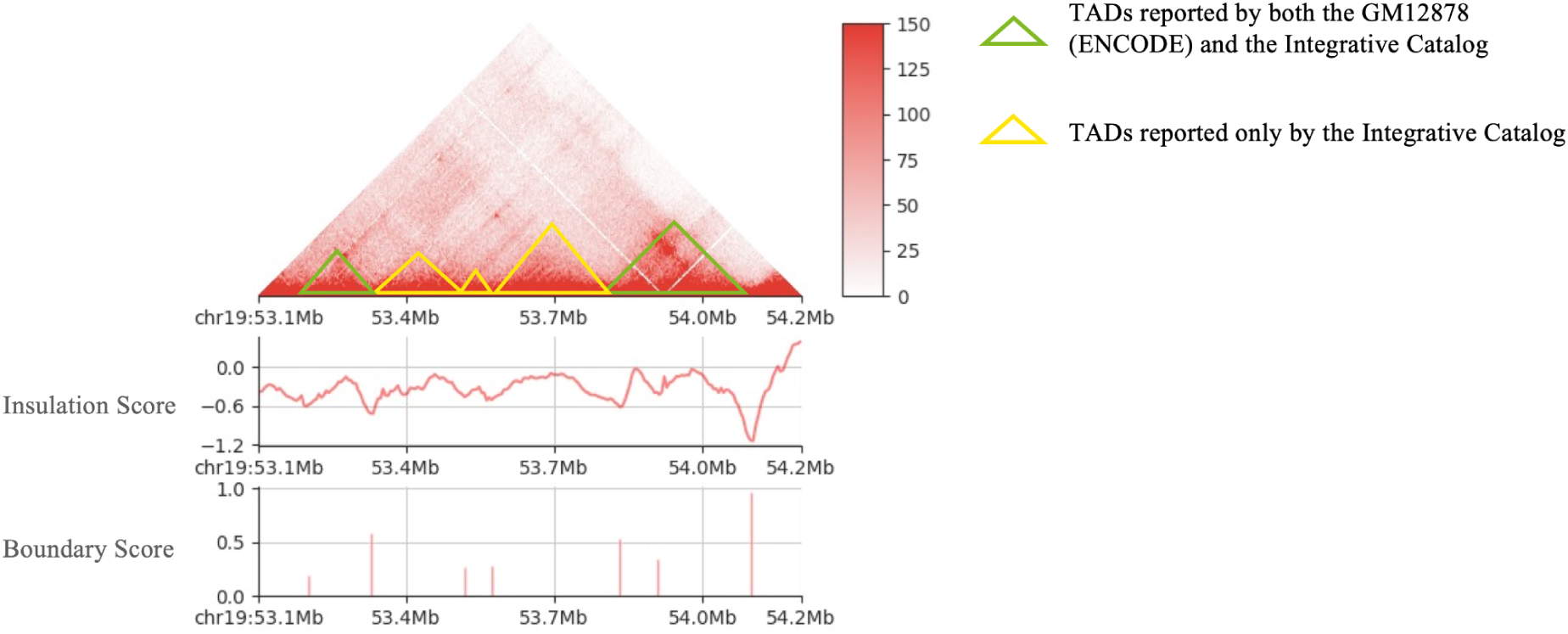
The comparison of TADs between GM12878 released by ENCODE and the Integrative TAD Catalog released in our study. **a).** The overlap of TADs between GM12878 released by ENCODE and the Integrative TAD Catalog generated using our customized pipeline in this study (one bp overlapped). Note that the 10,044 TADs of GM12878 released by ENCODE are at least one bp overlapped with 16,679 TADs in our Integrative Catalog (containing sub-TADs). Therefore, only 2,293 remaining TADs that were uniquely released in our Integrative Catalog are shown in the Venn diagram. **b).** Visualization of one region containing TADs that were identified in our Integrative Catalog but were missing in the GM12878 released by ENCODE. From top to bottom, these plots show the Hi-C contact maps, the insulation scores, and the corresponding boundaries with the boundary scores over this region. Green regions represent TADs identified by both GM12878 (ENCODE) and our Integrative Catalog, while yellow regions highlight TADs that were identified by our pipeline, not in GM12878 (ENCODE).

**Table 1.**
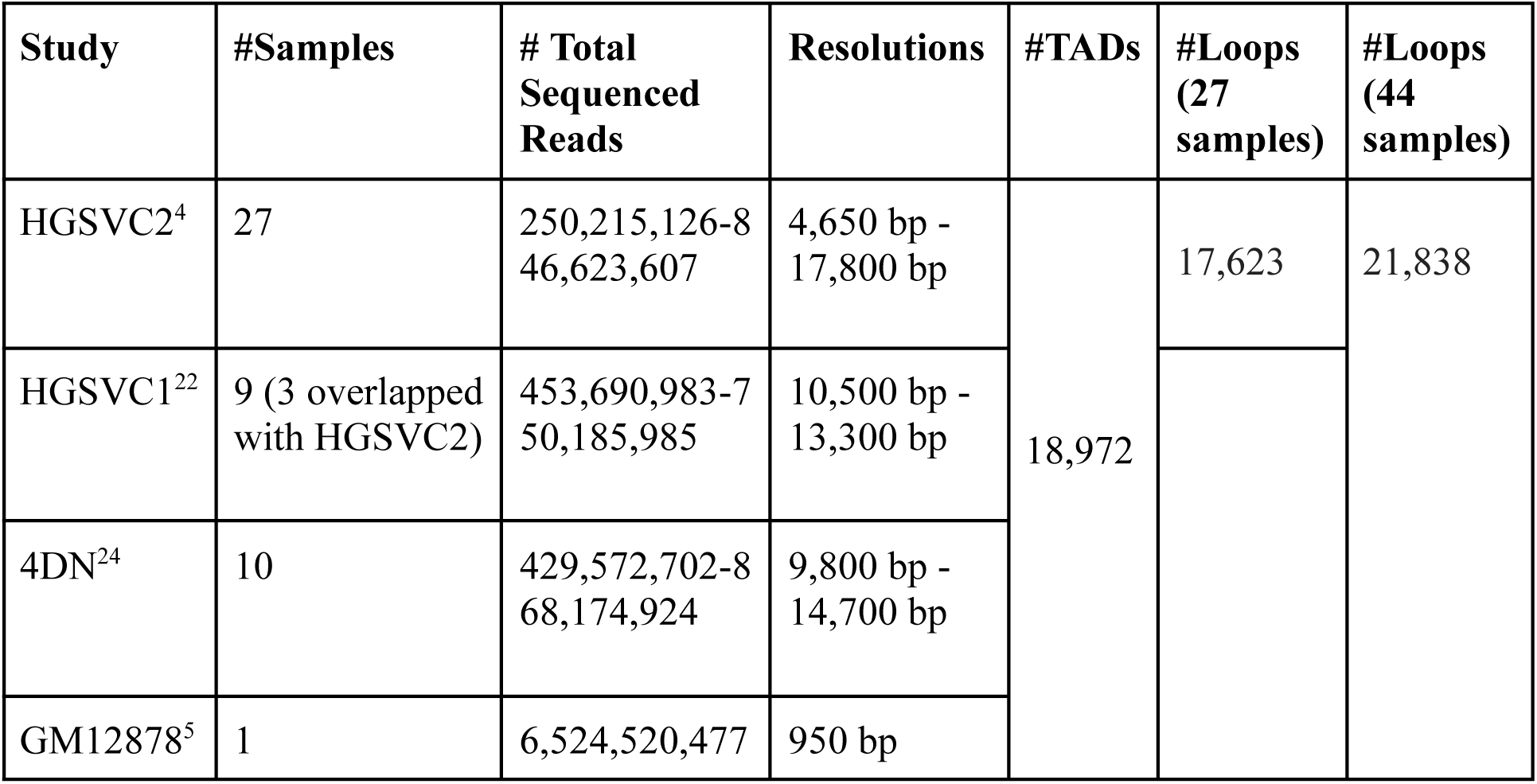
The Integrative Catalog in human 3D structure generated from this study aggregates Hi-C data from multiple projects. The Hi-C data from 44 individuals collected in this study come from HGSVC2^4^, HGSVC1^22^, 4DN project^22, 24^, and GM12878^5^ with the highest resolution. Three samples (HG00514, HG00733, NA19240) from HGSVC1 are re-sequenced in HGSVC2 with a higher resolution; thus, we kept those samples from HGSVC2 in our further analysis. The resolutions were calculated for each sample, and the TADs and loops were called on the 44 merged samples and 27 merged samples in HGSVC2, respectively.

| Study | #Samples | # Total Sequenced Reads | Resolutions | #TADs | #Loops (27 samples) | #Loops (44 samples) |
| --- | --- | --- | --- | --- | --- | --- |
| HGSVC2<sup>4</sup> | 27 | 250,215,126-8<br>46,623,607 | 4,650 bp -<br>17,800 bp | 18,972 | 17,623 | 21,838 |
| HGSVC1<sup>22</sup> | 9 (3 overlapped with HGSVC2) | 453,690,983-7<br>50,185,985 | 10,500 bp -<br>13,300 bp |  |  |  |
| 4DN<sup>24</sup> | 10 | 429,572,702-8<br>68,174,924 | 9,800 bp -<br>14,700 bp |  |  |  |
| GM12878<sup>5</sup> | 1 | 6,524,520,477 | 950 bp |  |  |  |

### 2. Identification of TAD boundaries for individuals in the HGSVC2 cohort

Next, we characterized and quantified TAD boundaries to explore how genetic variations like SVs disrupt TAD boundaries and bring about potential downstream changes in gene regulation and phenotypes (Table S6). 27 of the 44 individuals have been recently characterized by HGSVC2 with haplotypes containing genetic variation, including SVs^4^, which allowed us to detect the effect of SVs on 3D human genome chromatin conformation. We examined the map resolutions (compared with GM12878 in Rao’s study), read numbers, and contact numbers (Fig. 2) for each of the 27 samples, which inspired us to use the merging strategy in our Integrative Catalog TADs calling pipeline. In the end, we identified 14,612 boundaries (lenient set) and 5,884 strong boundaries (stringent set) in our Integrative Catalog with different boundary score cutoffs received from *IS* (Methods, Table S4, S5). For those 27 subjects in the HGSVC2 cohort, on average, we identified 12,410 boundaries (lenient set) and 5,299 strong boundaries (stringent set) (Fig. 3a, 3b). The HGSVC2 27 individuals come from five different super populations in Africa (AFR), East Asia (EAS), Europe (EUR), South Asia (SAS), and the Americas (AMR). We aggregated each individual by its super population and compared the numbers of TAD boundaries per individual genome (Fig. 3c, 3d). We did not observe any significant relationship between the different types of super populations and their insulation scores or boundary scores, which indicates that TAD features appear to be conserved across diverse human populations^16, 27^.

**Fig. 2.**
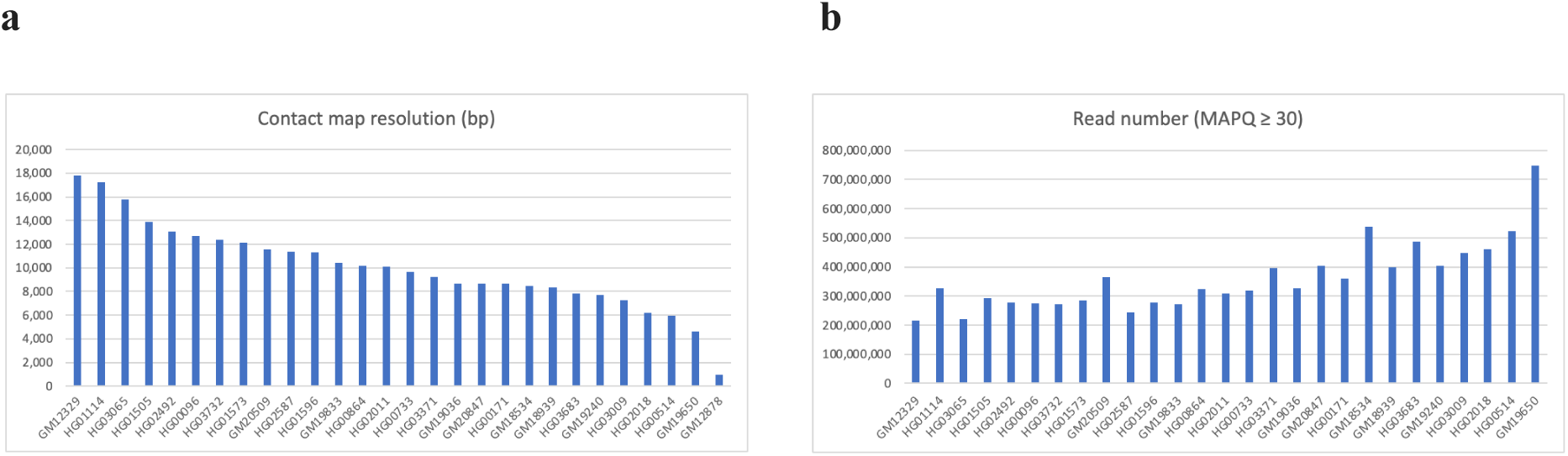
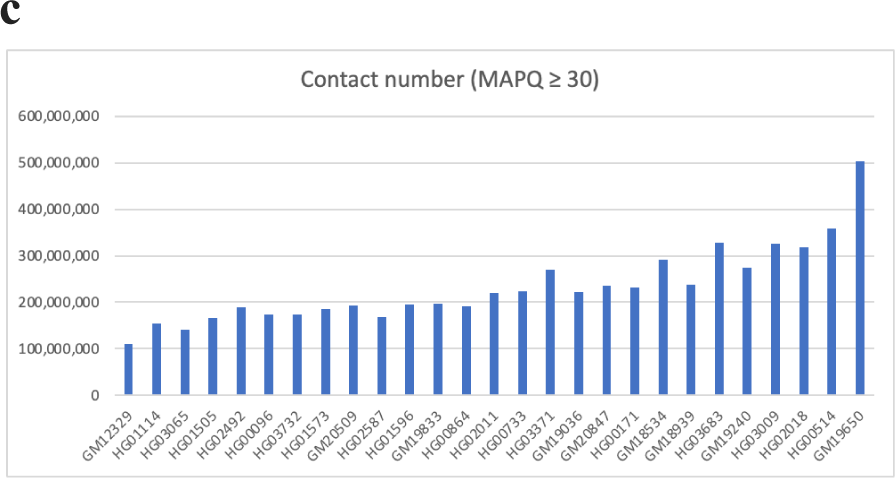
Map resolutions, read numbers, and contact numbers of each sample. **a).** The map resolutions of our 27 samples were compared with the GM12878 in the last columns. The x-axis represents the sample ID, from left to right, the resolution low to high; the y-axis shows the value of resolution in the base pair (bp). **b).** The number of read pairs of each sample with the filtered alignment quality based on the mapping quality score (MAPQ) ≥ 30. **c).** The number of Hi-C contacts of each sample with the filtered alignment quality for MAPQ ≥ 30.

**Fig. 3.**
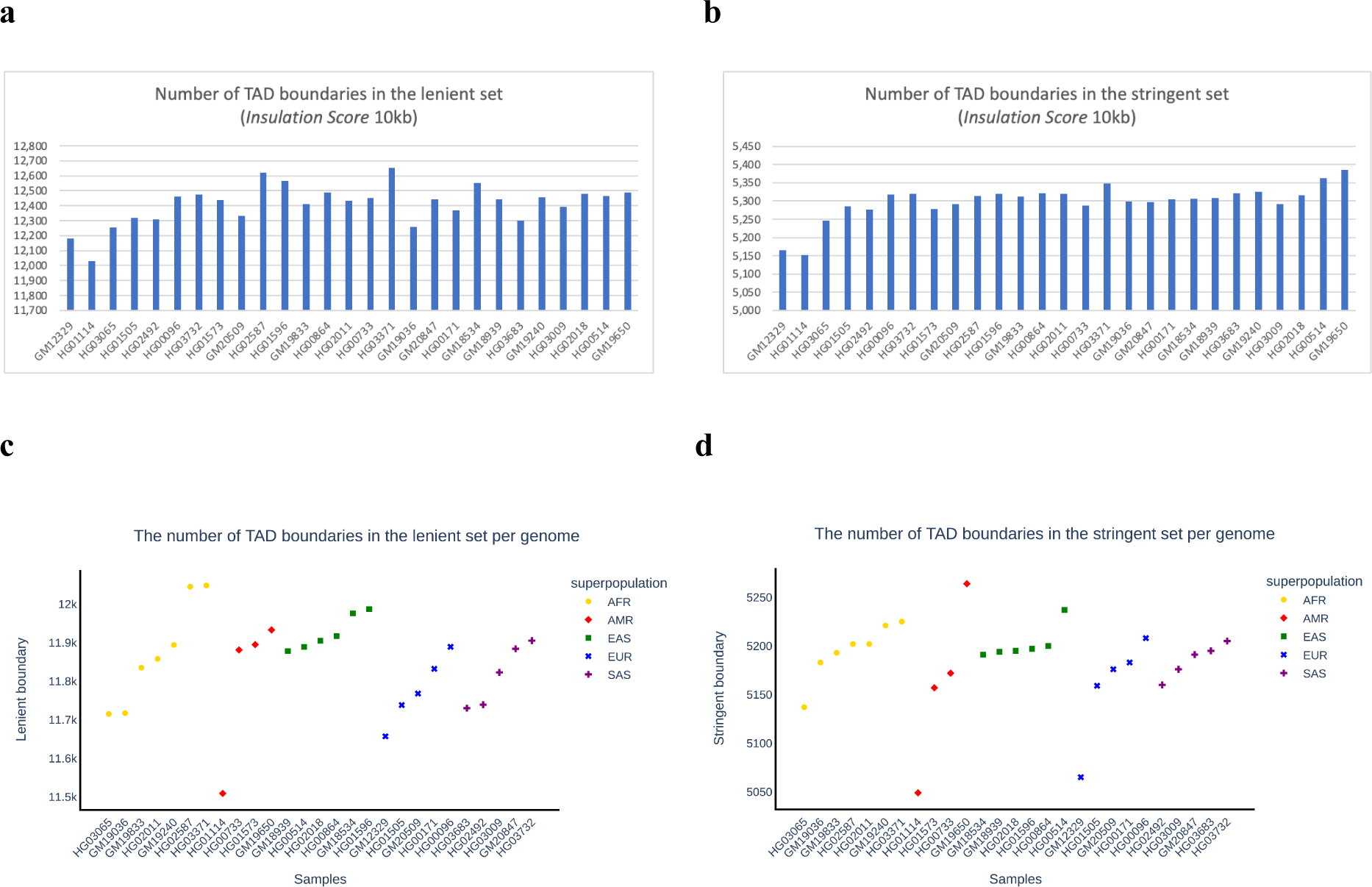
Distribution of TAD boundaries in 27 samples of HGSVC2. The x-axis represents sample IDs, with resolution ordered low to high from left to right in **a** and **b**, and with the super population ordered and represented by a different color as displayed in the color key in **c** and **d**. The y-axis shows the number of all TAD boundaries (lenient set in **a** and **c**), and that from the stringent set of TAD boundaries (stringent set, in **b** and **d**) detected with our pipeline in the 27 samples. All of these TAD boundaries were called with 10 kilobases (kb) resolution for each individual using *IS*.

### 3. SVs’ impact on 3D chromatin structure, gene expression, and splicing levels

We hypothesized that SVs in the human population disrupting TAD boundaries would have a functional impact on gene regulation^39^. To investigate the effects of SVs disrupting TAD boundary on the landscape of chromatin interaction, we intersected the identified TAD boundaries on the HGSVC2 samples with the SVs that are expression quantitative trait loci (SV-eQTLs) and splicing quantitative trait loci (SV-sQTLs) characterized in Ebert’s study^4^.

In total, we found 4,047 deletions and 5,512 insertions overlapped with the TAD boundaries identified in our Integrative Catalog. We used the boundary strength value (boundary score, BS) of each individual as a measurement of the disrupted chromatin structure to compare the BS changes between the homozygous genotype (0/0) for the SVs and other genotypes that contain at least one alternative allele (1/1, 0/1, and 1/0) for the SVs. BS is defined as the difference in the delta vector (the difference between the amount of insulation change 100 kb to the left and right of the central bin) between the nearest 5’ local maximum and 3’ local minimum relative to the boundary, which can be used to filter out the potential boundary^8^. Higher BS represents stronger boundaries, while lower BS refers to weaker boundaries. Significant changes in BS were observed in two deletions and four insertions (FDR *< 0.2*, Wilcoxon rank-sum test, two-sided) (Table S8). These findings suggest that removing or inserting particular TAD border sequences could alter the insulation between nearby TADs, thereby resulting in a change in the frequency of interactions between sequences in typically isolated domains.

We next examined if those SVs we found above that disrupt TAD boundaries have a potential effect on gene regulation using gene expression and splicing quantification from RNA sequencing (RNA-seq) of HGSVC2 samples. In the 26 HGSVC2 samples (the GM12329 sample was excluded due to a relatively low sequencing quality), we intersected the TAD boundaries with overlapped SVs that are also SV-eQTLs and SV-sQTLs (113 deletions and 120 insertions) and identified ten genes and 37 genes whose expression values were significantly changed for ten deletions and 32 insertions, respectively (FDR < 0.2, Wilcoxon rank-sum test, two-sided) (Table S9 and S10); We observed 173 splice junctions and 169 splice junctions whose splicing ratios were significantly changed for 71 deletions and 83 insertions, respectively (FDR < 0.205, Wilcoxon rank-sum test, two-sided) (Table S9 and S11). These results reveal that the deletion and insertion of TAD boundaries can result in significant changes in gene expression and splicing in human genomes.

By integrating these findings, our observations found one deletion (chr8-644401-DEL-5014) and one insertion (chr6-130293639-INS-295) within our 26 HGSVC2 call set disrupted regulatory sequences to influence local chromatin architecture and, in the meantime drive altered genes both on the expression and splicing levels. We observed that deletions tend to remove the TAD boundaries and fuse the adjacent TADs^33^, while insertion of sequence can split a single TAD into two adjacent TADs (neo-TAD)^57^ (Fig. 4).

**Fig. 4.**
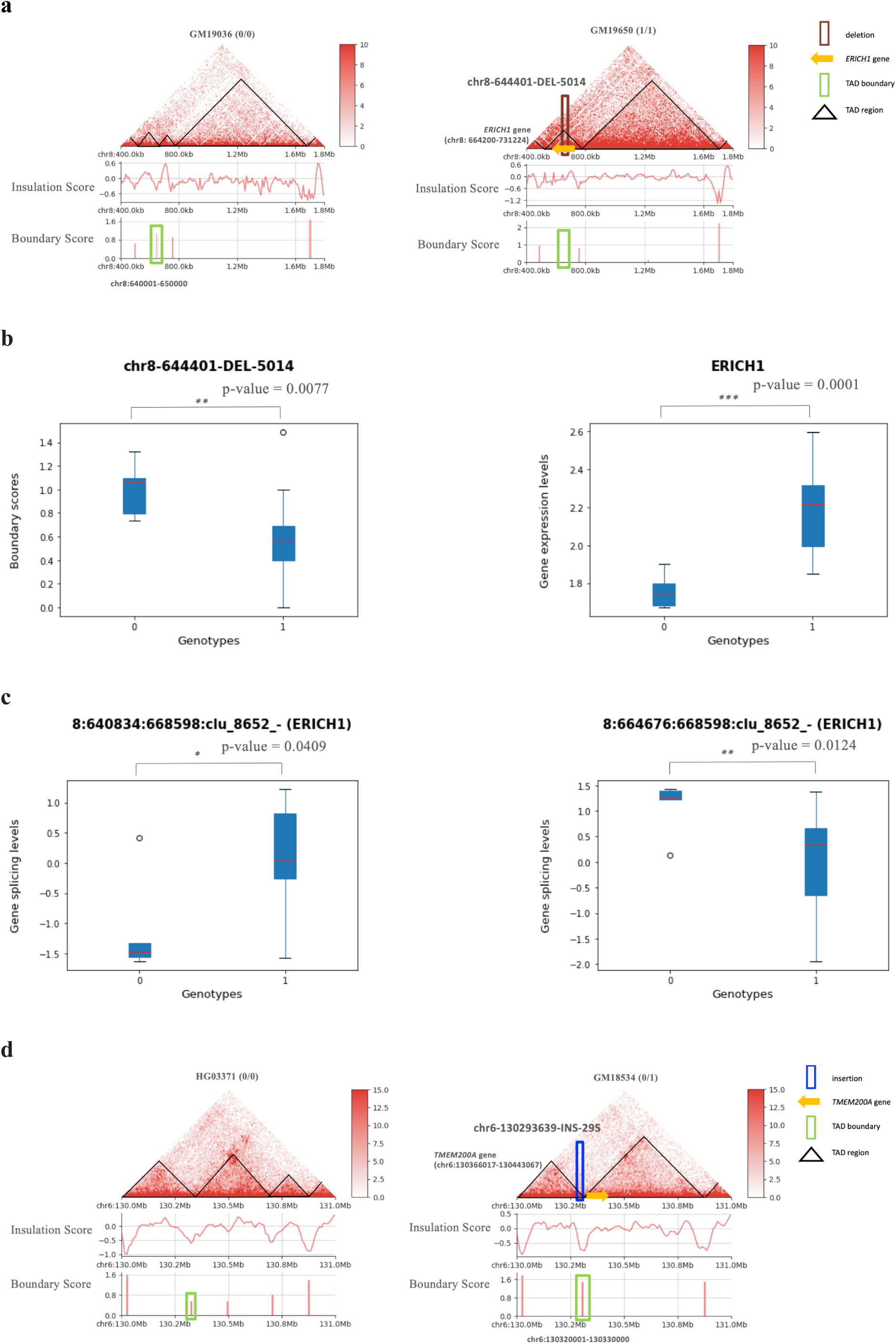
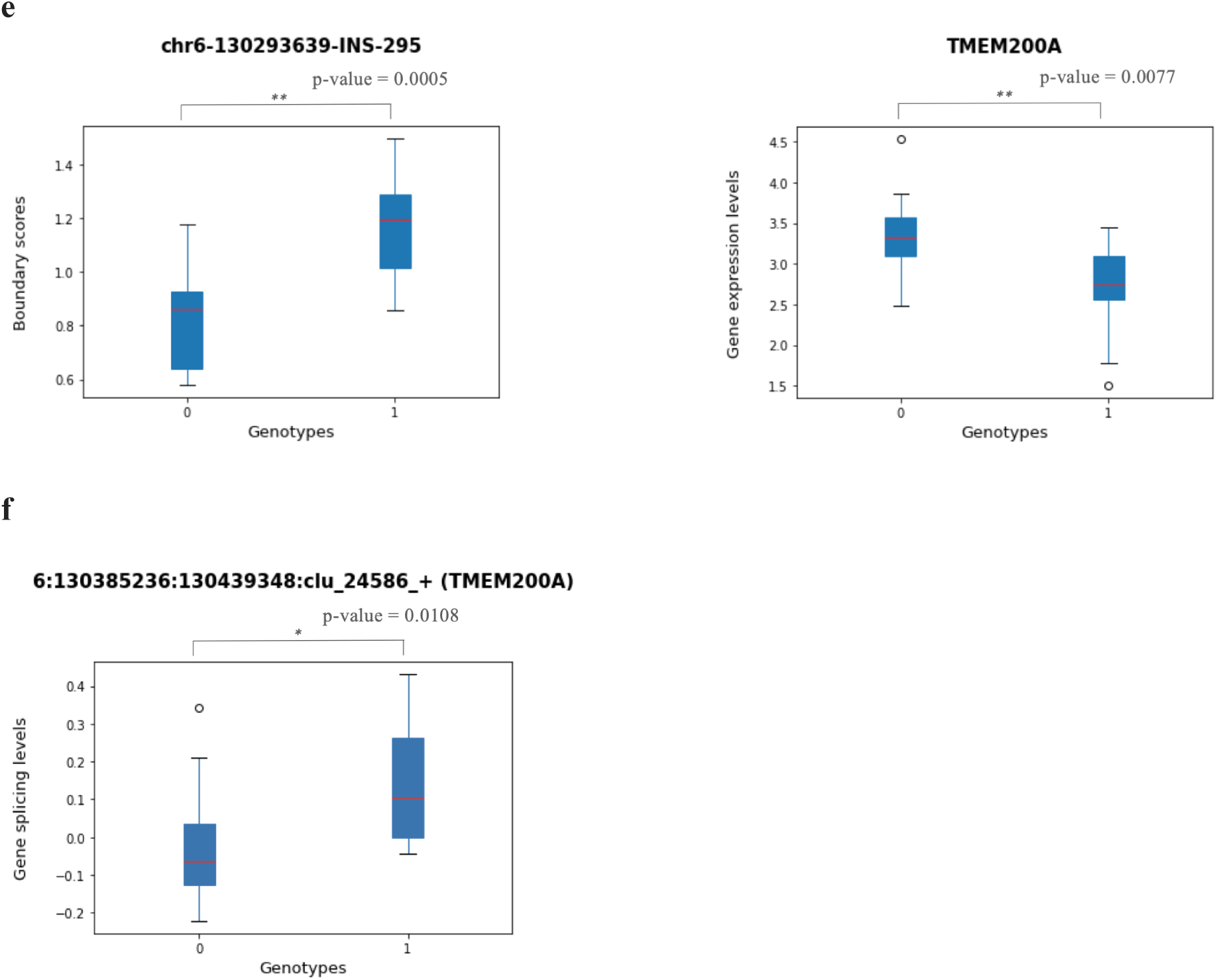
Visualization of two SVs that disrupt TAD boundaries with significant changes in boundary strength, gene expression, and splicing levels. **a).** A deletion (chr8-644401-DEL-5014) that disrupts the TAD boundary shows differences in Hi-C contact maps, boundary scores, and insulation scores for individuals with (genotype 1/1) and without (genotype 0/0) the deletion. The dark red rectangle represents the location of this deletion, and the blue arrow shows where the *ERICH1* gene is. The green rectangle highlights the TAD boundary’s location and corresponding boundary strength. The left figure is the GM19036 sample, whose genotype is 0/0, i.e., does not have this deletion, compared with the right sample GM19650, whose genotype is 1/1, i.e., carries this deletion. The boundary score panel shows that the right sample misses the TAD boundary where it carries that genomic deletion. **b).** Boxplots demonstrate the significant difference in gene expression for two different genotype categories, 0 (genotype 0/0) and 1 (genotypes 0/1, 1/0, or 1/1). The boxplot on the left shows the significant changes between the boundary score and different genotype categories of this deletion, while the right boxplot shows significant changes between the associated *ERICH1* gene expression values and different genotypes of this deletion. **c).** Boxplots show the significant difference between gene splice ratios (after log transformation and quantile normalization) for the splice junction clusters and genotypes of the same deletion. Specifically, we found two splicing quantitative trait loci QTLs (sQTLs) (8:640834:668598:clu_8652_- and 8:664676:668598:clu_8652_-) overlapped with this deletion, and both of which are associated with *ERICH1* gene. Subfigures **e** to **f** show the same comparisons for an example of insertion (chr6-130293639-INS-295) between the individual HG03371 (genotype 0/0) and GM18534 (genotype 0/1).

### 4. Catalog of chromatin loops across diverse individuals in human genomes

Besides TADs, we derived a comprehensive list of chromatin loops with the Hi-C datasets collected in this study^5^ (Table S6 and Table S14-S19). We constructed an integrated list of chromatin loops combining loops generated by two methods based on different overlap criteria. Chromatin loops of the 27 HGSVC2 merged set were predicted by both *HiCCUPS* GPU using the SCALE normalized matrix^5^ and *cooltools call-dots*^12^ which uses an iterative correction algorithm for matrix balancing (Imakaev et al. 2012 Nature Methods volume 9, pages 999–1003 (2012)). Loop calling was performed at both 5kb and 10 kb resolution. Our results for the merged loop set from these two models of loop calling show a very high consistency between HGSVC2 samples (Fig. 5a and 5b) as more than ∼74% of loops are exactly overlapped (17,623 loops; 74.0/75.6% *HiCCUPS*/*call-dots*) (Table S14, S15, and S16) and more than 80% loops are overlapped by adding 5 kb flanking length to the start and end site of the detected anchors (19,102 loops; 82.0/80.2% *HiCCUPS*/*call-dots*)) (Fig. 5). The addition of 4DN and HGSVC1 dilution Hi-C experiments to the HGSVC2 merged loop list led to a drop in the *HiCCUPS* and *call-dots* loop overlap (21,838 loops; 36.1/25.4% *HiCCUPS*/*call-dots*). *HiCCUPs* identified 60,494 merged loops, while *cooltools call-dots* identified 86,024 loops (Table S17 and S18). Overall, our integrated list of chromatin loops combining two loop calling methods amounts to 21,838 loops (Table S19). By adding a 5 kb flanking length to the start and end site of the detected anchors, 25,994 overlapped loops are found between *HiCCUPS* and *call-dots* (25,994 loops; 43.0/30.2% *HiCCUPS*/*call-dots*). The average distance between loop anchors was calculated for *HiCCUPS* and *call-dots* loops, with *call-dots* showing a slightly higher average distance between loops and wider bp distribution of the loops (Figure S5). Integration of the 44 Hi-C datasets has allowed us to generate a comprehensive chromatin loop catalog which we will use to study how genetic variation impacts specific chromatin contacts in the future.

**Fig. 5.**
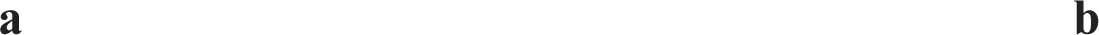
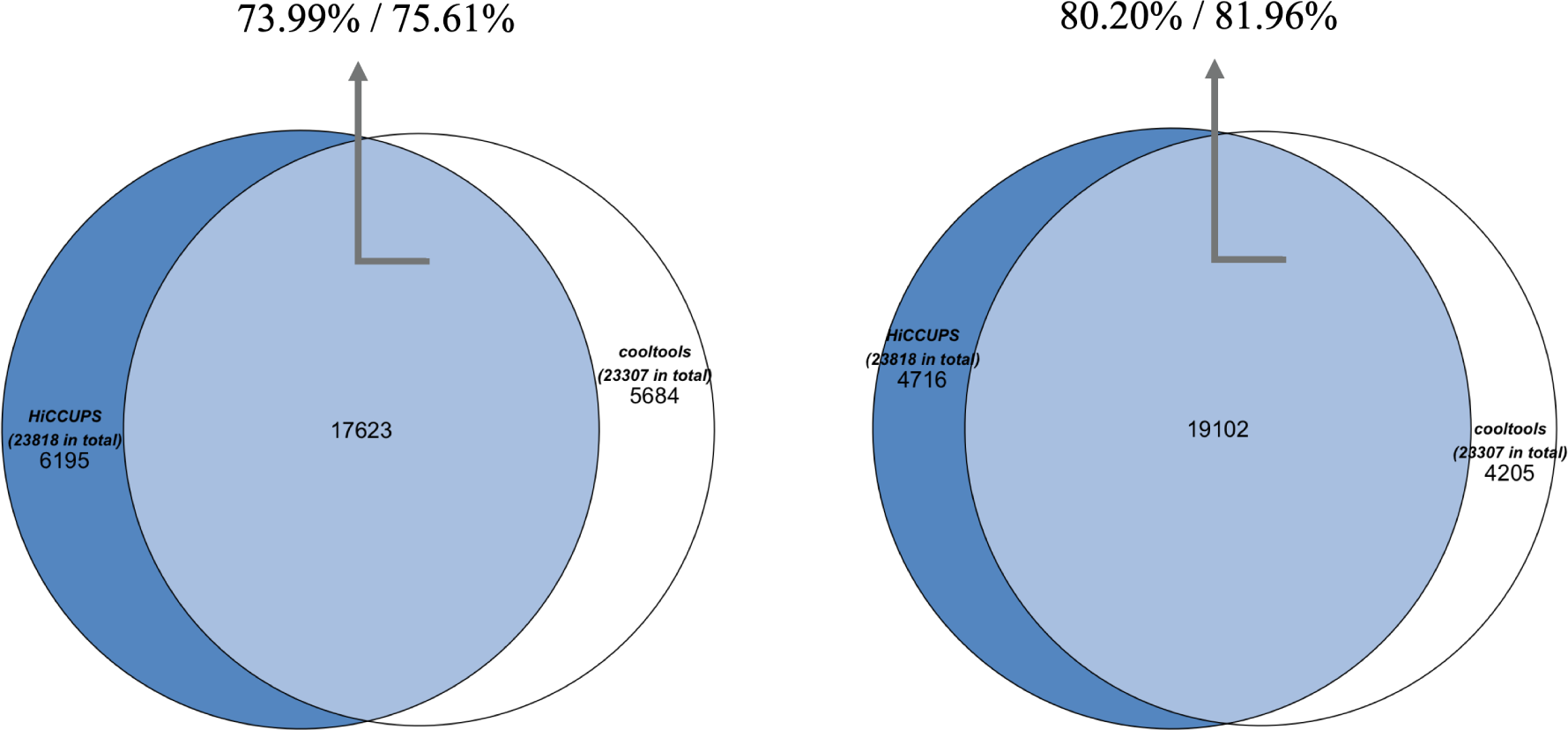
Comparison of loops detected by *HiCCUPs* and *cooltools* from the HGSVC2 merged Hi-C sample under 5kb and 10 kb merged resolutions. The agreement between *HiCCUPs* and *cooltools* loops is shown as Venn diagrams for our 27 merged Hi-C contact maps. The overlap between the two loop lists is colored in light blue, and the percentages of overlap with respect to each method are shown separately. **a)**. The exact same loop coordinates detected by both methods. **b).** The overlapped loop coordinates are produced by adding a 5 kb flanking region of the start and end site of anchors detected by both methods.

## Discussion

The 3D genome has revolutionized the field of genomics by providing a comprehensive view of the spatial organization of the genome. The availability of a comprehensive 3D genome map provides a valuable resource to assist with the understanding of human genome structure and function. Our study thus provides a comprehensive 3D genome catalog in diverse human genomes that serves as a valuable resource to elucidate chromatin organization in human genomes, by tackling current limitations of low resolution and lack of diversity in providing a map of 3D genome structure in healthy human individuals. To the best of our knowledge, our results provide the most comprehensive catalog of TADs and TAD boundaries in human LCLs (Table S2 to S6), which were produced by a customized Hi-C analysis pipeline that integrates Hi-C data generated from different platforms, with various sequencing resolutions aggregated from multiple data resources. We identified several significant examples of SVs that disrupt the genome 3D structure via genomic deletions or insertions in TAD boundaries, lead to gene regulation alterations, and also reported to contribute to pathological phenotypes such as developmental abnormalities, neurodegenerative disorders, and cancer^3, 13, 30–32, 77, 78^.

Previous research has revealed that alterations in the genome’s sequence can result in chromatin structural modifications, which might regulate gene expression and lead to various pathogenic phenotypes, such as aging-related disease, Alzheimer’s disease, cancers, and developmental disorders^3, 13, 35–38^. In recent studies, cancer development and progression have been proven to be influenced by allele-specific expression (ASE)^18, 19^. Thus, we sought to investigate if those two SVs and the associated eGenes and sGenes would have any potential contribution to the risk of complex diseases (Table S7). We found two of the genes (*ERICH1*, *TMEM200A*) were previously reported to be associated with various cancers (e.g., such as pancreatic cancer and gastric cancer) and tumor progression^15, 61–64^. Among them, *ERICH1* was reported to be related to the genomic imprinting regulatory mechanism (which is frequently linked to ASE^20, 21^) and differential allelic expression^17, 65^. Interestingly, one recent study reported that the expression of *TMEM200A* was notably high in gastric cancer, and overexpression of *TMEM200A* would shorten the overall survival of gastric cancer patients^63^. They concluded that upregulated *TMEM200A* would serve as a promising prognostic marker for gastric cancer and is closely associated with the tumor microenvironment.

However, there is room for improvement in our current analysis to note here. As more Hi-C data is available (e.g., HGSVC3 Hi-C sequencing data), there is an essential need for us to add those data to our current Integrative Catalog, and our pipeline is readily available to support such analysis. Another limitation in our analysis is that we only investigated SVs in the current HGSVC2 eQTL/sQTL results (deletions and insertions)^4^. We will extend our investigation of the impact of other types of SVs (e.g., duplications, inversions) on the chromatin structure and gene regulation in the future. In addition, the Telomere-to-Telomere (T2T) Consortium has released the first complete and gapless sequence of the human genome sequence, named T2T-CHM13^55^, which will complement the standard human reference genome that was used in our study, known as Genome Reference Consortium build 38 (GRCh38). With this new reference genome available, we are able to map our Hi-C reads to T2T and potentially personal assemblies with no gaps and unprecedented accuracy.

We envision that the comprehensive catalog of TADs, TAD boundaries, and chromatin loops produced from this study will provide a reference map of known 3D genome structures of human genomes. Our findings have critical implications for understanding human whole-genome sequencing data in disease diagnosis and precision medicine. Genomic variation, including all types of SVs that overlap or intersect with TAD boundaries in human genomes, should be further investigated to better understand the molecular basis of human genetic diseases^26^.

In summary, we generated a high-resolution 3D genome profile of TADs, TAD boundaries, and chromatin loops of diverse human genomes. This comprehensive catalog allows us to associate SVs with 3D genome structure to reveal the importance of TAD boundary sequences for genome function and regulatory mechanisms. Our study provides insight into the significance of integrative analysis of SVs and TADs, in that deletions and insertions overlapping with and proximal to TAD boundaries should be carefully examined in disease studies. We believe that the released Integrative Catalog of TADs and TAD boundaries in the human genome will assist the investigation of genomic variation, gene regulation, 3D genome structure, and their impact on and association with human diseases, with promises to provide insights into disease etiology and therapeutic discovery^41^.

## STAR Methods

### Resource availability Lead contact

Further information and request for resources and reagents should be directed to and will be fulfilled by the lead contact, Xinghua Shi.

### Materials availability

This study did not generate new unique reagents.

## Method details

### Hi-C data collection

We have collected Hi-C data on 44 samples from HGSVC2, HGSVC1, 4DN, and GM12878 (Table S12). The first dataset (HGSVC2) is the Hi-C data for the 27 HGSVC2 samples generated from the Human Genome Structural Variation Consortium. Hi-C libraries were generated with 1.5 M human cells as input using Proximo Hi-C kits v4.0 (Phase Genomics, Seattle, WA) according to the manufacturer’s protocol with the following change: cells were crosslinked, quenched, lysed sequentially with Lysis Buffers 1 and 2, and liberated chromatin immobilized on magnetic recovery beads. During the fragmentation process, a cocktail of 4-cutter restriction enzymes (DpnII (GATC), DdeI (CTNAG), HinfI (GANTC), and MseI (TTAA) was utilized to enhance coverage and facilitate haplotype phasing. Fragmented chromatin was proximity ligated for 4 hours at 25°C after fragmentation and fill-in with biotinylated nucleotides. The crosslinks were then reversed. Magnetic streptavidin beads were used to purify DNA and retrieve biotinylated junctions. The dual-unique indexed library was created using bead-bound proximity ligated fragments and Illumina sequencing chemistry. Fluorescent-based assays, such as qPCR with the Universal KAPA Library Quantification Kit and Tapestation, were used to assess the Hi-C libraries (Agilent). The libraries were sequenced at New York Genome Center (NYGC) in a paired-end 150 bp format on an Illumina Novaseq 6000.

The second (HGSVC1) and third (4DN) pilot datasets are the Hi-C sequencing data which has been generated for the 1000 Genomes Project SV group by the laboratory of Bing Ren using the Illumina HiSeq (2000, 2500 or 3000) paired-end sequencing^22, 24, 54^ and were performed using a “dilution” 6-cutter HindIII (AAGCTT) protocol. The second pilot dataset came from Hi-C sequencing for three trios: Yoruba (NA19238, NA19239, NA19240), Han Chinese (HG00512, HG00513, HG00514), and Puerto Rican (HG00731, HG00732, HG00733). The HG00514, HG00733, and NA19240 were excluded from this dataset since these samples are re-sequenced in the HGSVC2 study. For the same reason, the GM19238 from the third (4DN) pilot dataset was also excluded.

We also included the Hi-C data from GM12878 B-lymphoblastoid cells in our analysis from Rao et al., which has almost 5 billion mapped paired-end read pairs and is considered the largest coverage for the accurate identification of 3D chromatin structures. The sequence data were produced as in Rao et al. (2014)^5^ and can be downloaded from ENCODE.

### Hi-C Data Processing

We required a minimum alignment quality for each read included in our Hi-C maps. We used the mapping quality score (MAPQ), which quantifies the probability that a read is misplaced, to filter out the read pairs where the alignment of one or both reads fails to meet these two thresholds: MAPQ > 0 and MAPQ ≥ 30 (Fig. 2b)^5^. In our study, we used the MAPQ ≥ 30 filtered data to be stringent about avoiding false positives caused by poor alignments. The result of filtering low-quality alignments is a list of Hi-C contacts (Fig. 2c).

For each sample, the raw resulting reads were mapped to the GRCh38 reference genome and processed using *Juicer* software tools (version 1.6) with default aligner BWA mem^7, 73^. Unmapped reads such as abnormal split reads and duplicate reads were removed, and low mapping quality read pairs were filtered out if their mapping quality value MAPQ was less than 30. Those filtered read pairs (Hi-C contacts) were subsequently used to construct chromatin contact maps for each sample by *Juicer*. To build a Hi-C contact map on an Integrative Catalog of LCLs basis, contacts were pooled across all 44 individuals using the *mega.sh* script provided in *Juicer*. The outputs of the previous processes are two sets of *.hic* files, a special format of the highly compressed binary file to store contact matrices with various resolutions (i.e., bin sizes) and can be mainly supported by *Juicer*, *JuicerTools*, and *Juicebox* command line tools for downstream analysis and visualization^9^. All analyses and results reported in our study employ the contact frequency matrices normalized with SCALE matrix balancing^7^, which remedies the issue that sometimes KR normalization does not provide coverage for a particular region or chromosome^23^.

### Calculation of Hi-C map resolution

To decide the applicable TAD map resolution to be called for each sample, we first applied the script from *Juicer* to calculate the map resolution of each sample^7^. The map resolution is intended to reflect the finest scale at which local features can be reliably detected. The lowest resolution is 17,800 bp (GM12329), while the highest is 4,650 bp (GM19650). *Juicer* created a specific .*hic* file format to describe the input reads under nine different bin sizes (base-pair-delimited resolutions: 2500000, 1000000, 500000, 250000, 100000, 50000, 25000, 10000, 5000), and usually based on the sequencing depth of the Hi-C file, 5 kb or 10 kb resolution will be used to call *Arrowhead*^7^. Considering that GM12878, with the largest contact map and deepest depth of sequencing in its Hi-C file up to now (map resolution of 950 bp), was called under 5 kb resolution in the *Arrowhead*^5^, we chose the same 5 kb as the practical resolution of TAD calling on all of our 44 merged samples (kilobase map resolution of 200 bp).

We applied the method used by Rao et al.^5^ to calculate the Hi-C map resolution of our 27 samples (Fig. 2a). The map resolution is defined as the smallest bin size that 80% of bins have at least 1,000 contacts, which is intended to reflect the finest scale at which local features can be detected reliably. The scripts for calculating the Hi-C resolution of each sample were directly downloaded from Rao’s study^5^.

### Identification and visualization of TAD and TAD boundaries

Two TAD callers, *JuicerTools* (version 1.22.01)’ *Arrowhead* and *Insulation Score* (*IS*), were used and compared to call TADs for 43 (except GM12878) samples respectively at 10 kb resolution from the human lymphoblast line^7, 8^. *Arrowhead* was more accurate and sensitive for ultra-high resolution data and focused on detecting the corners of the domains to locate the boundaries of TADs, while *IS* algorithm was initially created to find TAD boundaries on Hi-C data with a relatively low resolution^25^. For pool calling, SCALE normalized merged contact matrix (44 samples) at 5 kb resolution was used to calculate the insulation score and corresponding boundary score (BS) using the *FAN-C* toolkit version 0.9.26b2 with default parameters to detect the TAD boundaries at a 100 kb window size (which was referenced from the 4DN domain calling protocol)^10, 24^. The same method was applied for sample calling and used the SCALE normalized contact matrix at 10 kb of each sample as input. The *IS* defines a sliding window and adds up contacts in this window to align the Hi-C matrix diagonal. The regions with low insulation scores (high boundary scores) are insulating and referred to as TAD boundaries, and regions with high insulation scores (low boundary scores) are most typically found inside domains and referred to as TAD regions, which were also considered as the regions between the two neighbored TAD boundaries^10^.

The sex chromosomes X and Y were eliminated from all analyses because of the gender disparities in our samples. ENCODE has processed the GM12878 files for hg38 and released the TADs called by *Arrowhead* on the SCALE normalized Hi-C matrices^60^. Thus, we compared their TAD results with the GM12878 result processed by our pipeline to determine a minimum boundary strength cut-off value (0.17) for GM12878, which gave the most repetitive results. Comparing our minimum boundary score cut-off value of 0.17 with the 0.2 value used in the 4DN TAD calling protocol, we simply decreased 0.03 to the 4DN strong boundary score cut-off value of 0.5 as our strong cut-off value at 0.47. Those values were then transformed to a percentile to find the corresponding cut-off scores in our LCLs call set. After the removal of the missing and duplicated boundary scores, a minimum boundary score cut-off value of 0.1880268762332596 and a strong boundary score cut-off value of 0.5477218522527745 were chosen. The insulation scores, TAD boundaries, and corresponding boundary scores were visualized using *Juicebox* software and the *FAN-C* toolkit in Python 3.7^9, 10^.

We next sought to generate the most comprehensive TADs call set aligned with our TAD boundary locations for more downstream analysis (Fig. 6). To be specific, 1) we first called the TADs directly from *Arrowhead* under 5 kb resolution to be our TADs callset1; 2) the boundaries we previously received from *IS* were first converted to TADs and then be filtered by a) excluding those TADs that contain more than ten percent of the length of those TADs that have missing insulation scores and b) removing TADs that don’t have any maxima of insulation scores to be our TADs callset2; 3) We then used “*bedtools intersect*”^71^ to find the overlapped regions that are both detected by *Arrowhead* (TADs callset1) and *IS* (TADs callset2) when the reciprocal overlap was above 50%. For those overlapped regions that contain more than one TAD detected from *Arrowhead*, we kept the TAD locations identified from *Arrowhead* instead of *IS* since *Arrowhead* is able to find the sub-TADs nested or overlapped within the larger TADs, which are demonstrated to have more cell or species-specific gene regulation activities within it^5, 16, 56^. We subtracted those regions both from callset1 and callset2 to avoid duplicates; 4) We then merged the remaining overlapped regions from step 3 by the smallest start position and largest end position for each TAD; 5) We finally added the TADs that distinctly detected from *Arrowhead* and *IS*, the sub-TAD regions from step 3, the merged regions from step 4 to generate our final comprehensive TAD catalog.

**Fig. 6.**
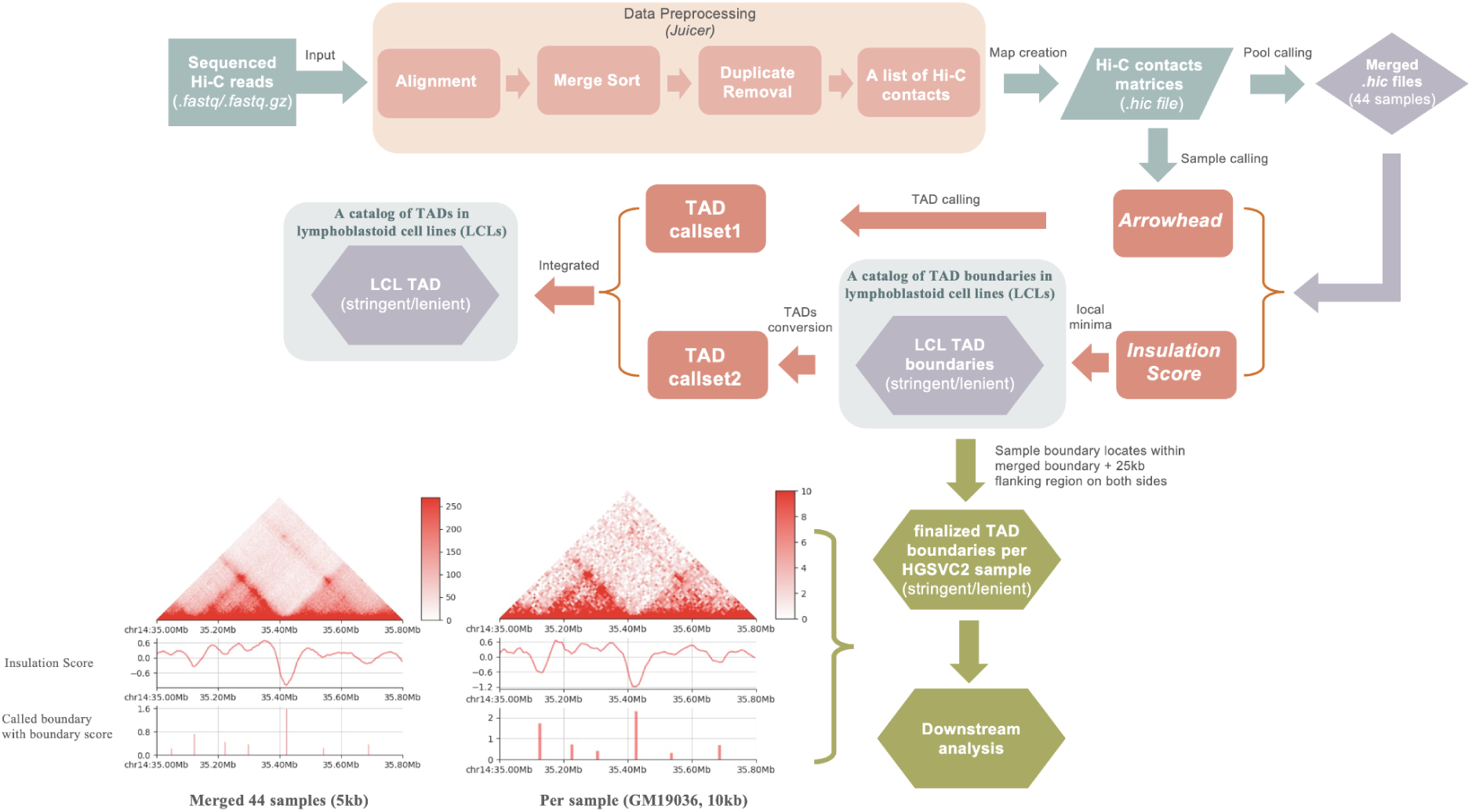
The step-by-step workflow to process raw Hi-C data into TADs and TAD boundaries in our Hi-C analysis pipeline. 44 samples’ raw reads files were used as input in *Juicer* to preprocess and create Hi-C maps, which were binned at multiple resolutions. *Insulation score* algorithms were applied to call an initial TAD boundary for each sample. All 44 *.hic* files were then merged together to create a “mega” map and used as an input of *Arrowhead* and *Insulation Score* algorithms to call TADs, and TAD boundaries for the LCLs merged call set. A finalized TAD boundary results for each individual were defined as those sample boundaries located within the merged boundary plus 25 kb flanking regions on the left side of the boundary start site and the right side of the boundary end site^11^. The two figures located in the bottom left corner are shown as a comparison between the merged subjects level and single subject level, which includes the Hi-C contact maps, the insulation scores, and the boundary strengths for the merged call set (5kb) and the GM19036 (10kb) sample over the region chr14: 35Mb-35.8Mb.

### Individual genotype verification

We used *verifyBamID* to verify whether the Hi-C sequencing reads in our aligned files match with the PanGenie genotyped SV calls of the 44 individuals used in this study^67, 68^. It can also identify whether the reads have been contaminated or swapped by a mixture of two samples. We did this for each of our samples (Table S13), and we observed that one of the samples (GM19204) has 12% or more of non-reference based observed in reference cites, which is much greater than a normal 2% standard criteria and indicated that this sample is very likely to have contamination. For this reason, we ensured to exclude this sample from any of our genotype-related downstream analyses for accuracy.

### SV-eQTLs and SV-sQTLs candidates selection

Gene expression and splicing levels’ quantifications of HGSVC2 26 samples (excluded GM12329) were extracted from RNA-seq and produced as previously described in our HGSVC2 study^4^. SV-eQTLs, SV-sQTLs results, and the PanGenie genotyped SV calls used in this study were obtained on the same 26 samples from the HGSVC2 project^4^. The SV-eQTLs result includes 34,745 deletions and 25,516 insertions; the SV-sQTLs result contains 44,945 deletions and 33,950 insertions. Only deletion and insertion alleles that: 1) were genotyped in our pangenie SVs calls; 2) were present in at least five samples; and 3) the homozygous for the reference allele of SVs (0/0) were present at least once in our sample of 26 individuals were kept in our final analysis.

### Quantification of the effects of candidate SVs on TAD boundary

A Wilcoxon rank-sum test (Mann–Whitney U test) was performed on each of those SVs located within the flanking TAD boundaries between the homozygous reference (0/0) and the heterozygous/homozygous deletions (1/1, 0/1, and 1/0) for the 26 samples using the boundary scores calculated by *IS*, and an FDR < 0.2 was considered as significant. If multiple TAD boundaries per sample were located inside one single TAD boundary from the merged call set, we used the median value of those samples’ TAD boundaries. All test statistics were estimated using the Scipy library, and box plots were generated using Matplotlib in Python 3.7.3. The False discovery rate (FDR) correction was conducted in the *qvalue* 2.28.0 package in R 4.2.1.

### HGSVC chromatin loop calling

The chromatin loops for the merged set and individual samples were identified by *HiCCUPS* GPU^5^ at 5 kb and 10 kb resolution in a merged resolution manner on the SCALE normalized Hi-C matrix with all other parameters at default. For comparison, an alternative modular Hi-C analysis pipeline was used for analyzing the individual Hi-C samples, called *distiller-nf*^72^. An iterative correction algorithm for matrix balancing, *cooler balance,* was used to balance *.cool* matrix files generated by distiller^69^. The *cooltools* Hi-C data analysis package has a loop-calling function named call-dots^12^. *Call-dots* is a re-implantation of the *HiCCUPS* loop-calling algorithm and was used as an additional model for loop calling of all individual Hi-C samples. For a generation of the merged loop list, *HiCCUPS* was used to generate a merged loop list on the merged *.hic* file under 5 kb and 10 kb merged resolutions. The merged *cooltools* loop list was generated by converting the *.hic* file to a .*cool* file using hic2cool convert^74^. The *cooler* support library function, cooler balance, was used for matrix balancing^69^. Dots were called at 5kb and 10kb resolution with the following parameters: fdr = 0.1, tile_size = 20000000, max-nans-tolerated = 7, kernel-width =7, kernel-peak =4, dot-clustering-radius = 20,000. 5kb and 10kb resolution loop lists were merged using *HiCExplorer* tools’ hicMergeLoops^70, 75^. *Bedtools* pairtopair^71^ was used to identify loop overlap between *HiCCUPS* and *call-dots.* The final reported loop lists included in this study were detected using both models, and only those detected by both models were reported to reduce the false-positive loop results (in a stringent and robust overlapped criteria manners introduced in the results section).

## Data availability

The raw sequencing Hi-C data generated by HGSVC2 discussed in this publication can be downloaded directly at the following link: (http://ftp.1000genomes.ebi.ac.uk/vol1/ftp/data_collections/HGSVC2/working/20200512_Hi-C/)^4^. The raw sequencing Hi-C data of HGSVC1 generated by Bing Ren can be found at (http://www.ebi.ac.uk/ena/data/view/PRJEB11418). The raw sequencing Hi-C and other processed data files generated by Ren lab are available through the 4D Nucleome data portal (https://data.4dnucleome.org/publications/b8c7c5f5-c76f-457f-9a0d-6c567924b816/#expsets-table). The raw sequencing GM12878 Hi-C data have been deposited into NCBI’s Gene Expression Omnibus (GEO) accession GSE63525^5^, as well as through the Encode project portal (https://www.encodeproject.org/experiments/ENCSR410MDC/). The RNA-seq data discussed in this study have been deposited in the following link: (http://ftp.1000genomes.ebi.ac.uk/vol1/ftp/data_collections/HGSVC2/working/20200627_RNAseq_JAX/). The PanGenie genotyped SV calls can be downloaded directly at the following link (http://ftp.1000genomes.ebi.ac.uk/vol1/ftp/data_collections/HGSVC2/release/v2.0/PanGenie_results/)4. The eQTL calls can be downloaded from the https://drive.google.com/file/d/1S0nARNKakaQ6vAdRr0dsMIv3_dZzD54E/view?usp=share_link, and the sQTL calls can be downloaded from the https://drive.google.com/file/d/1L6KhNn-RkC2GA0XrJmmqNbk2urNCH9JZ/view?usp=share_link. On reasonable request, the respective authors can provide additional data to support the conclusions of this study.

## Data and code availability

All data reported in this paper will be shared by the lead contact upon request.

All original code for the statistical analysis and pipeline has been deposited on GitHub https://github.com/caragraduate/Hi-C-integrative-catalog and is publicly available as of the date of publication.

Any additional information required to reanalyze the data reported in this paper is available from the lead contact upon request.

## Acknowledgment

This research is supported by National Institutes of Health (NIH) grants U24HG007497 (to C.L., E.E.E., J.O.K., T.M.), U01HG010973 (to T.M., E.E.E., and J.O.K.), and R01HG002385 and R01HG010169 (to E.E.E.); the German Federal Ministry for Research and Education (BMBF 031L0184 to J.O.K. and T.M.); the German Research Foundation (DFG 391137747 to T.M.); the German Human Genome-Phenome Archive (DFG [NFDI 1/1] to J.O.K.); the European Research Council (ERC Consolidator grant 773026 to J.O.K.); E.E.E. is an investigator of the Howard Hughes Medical Institute.

We want to thank Michael Zody, Michael Talkowski, Mark Chaisson, and Weichen Zhou from HGSVC Functional Analysis Working Group for providing critical and constructive advice and discussion about Hi-C analysis. We thank Mohammad Erfan Mowlaei and Emily Thyrum for their feedback on the project. We sincerely extend our gratitude to the people who contributed samples to the 1000 Genomes Project.

## Author contributions

The study was designed by C.Li. and overseen by X.S., HGSVC (co-chaired by T.B., J.O.K., E.E.E., and C.L.), and the HGSVC Functional Analysis Working Group. C.Li. conducted data analysis, statistical analysis, and visualization of the Hi-C and associated data and results.

M.J.B. assisted in the analysis of Hi-C data. C.Li. and S.S. performed loop calling. C.Li. and X.S. wrote the manuscript, incorporating contributions from all other co-authors. All authors read and approved the final manuscript.

## Consortia

The members of the Human Genome Structural Variation Consortium (HGSVC) are Haley J. Abel, Hufsah Ashraf, Peter A. Audano, Anna O. Basile, Christine Beck, Marc Jan Bonder, Harrison Brand, Marta Byrska-Bishop, Mark J.P. Chaisson, Yu Chen, Ken Chen, Zechen Chong, Nelson T. Chuang, Wayne E. Clarke, André Corvelo, Scott E. Devine, Peter Ebert, Jana Ebler, Evan E. Eichler, Uday S. Evani, Susan Fairley, Paul Flicek, Mark B. Gerstein, Maryam Ghareghani, Ira M. Hall, Pille Hallast, William T. Harvey, Patrick Hasenfeld, Alex R. Hastie, Wolfram Höps, PingHsun Hsieh, Sarah Hunt, Miriam K. Konkel, Jan O. Korbel, Sushant Kumar, Charles Lee, Alexandra P. Lewis, Chong Li, Bin Li, Yang I. Li, Jiadong Lin, Mark Loftus, Tsung-Yu Lu, Rebecca Serra Mari, Tobias Marschall, Ryan E. Mills, Zepeng Mu, Katherine M. Munson, David Porubsky, Benjamin Raeder, Tobias Rausch, Allison A. Regier, Jingwen Ren, Bernardo Rodriguez-Martin, Ashley D. Sanders, Martin Santamarina, Xinghua Shi, Chen Song, Oliver Stegle, Michael E. Talkowski, Luke J. Tallon, Jose M.C. Tubio, Aaron M. Wenger, Xiaofei Yang, Kai Ye, Feyza Yilmaz, Xuefang Zhao, Weichen Zhou, Qihui Zhu, and Michael C. Zody.

## HGSVC Functional Analysis Working Group

The members of the Human Genome Structural Variation Consortium (HGSVC) functional analysis working group are Anna O. Basile, Bernardo Rodriguez-Martin, Chong Li, Marc Jan Bonder, Marta Byrska-Bishop, Mark J.P. Chaisson, Ken Chen, Wolfram Höps, Yang I. Li, Ryan E. Mills, Zepeng Mu, Sabriya Syed, Qingnan Liang, Michael E. Talkowski, Yukun Tan, Matthew Jensen, Weichen Zhou, and Michael C. Zody.

